# Precise identification of cancer cells from allelic imbalances in single cell transcriptomes

**DOI:** 10.1101/2021.11.25.469995

**Authors:** Mi K. Trinh, Clarissa N. Pacyna, Gerda Kildisiute, Nathaniel D. Anderson, Eleonora Khabirova, Sam Behjati, Matthew D. Young

## Abstract

A fundamental step of tumour single cell mRNA analysis is separating cancer and non-cancer cells. We show that the common approach to separation, using shifts in average expression, can lead to erroneous biological conclusions. By contrast, allelic imbalances representing copy number changes directly detect the cancer genotype and accurately separate cancer from non-cancer cells. Our findings provide a definitive approach to identifying cancer cells from single cell mRNA sequencing data.

---

Single cell mRNA sequencing has enabled transcriptomic profiling of tumours and their environment with data being generated across the entire spectrum of human cancer. Studying the cancer transcriptomes depends on accurate identification of cancer cells. Therefore, the foundational step of tumour single cell analyses is separating cancer from non-cancer cells.

Expression of cancer specific marker genes can identify cancer cells in some cases, but is generally insufficiently precise, especially without corroboration from readouts such as cellular morphology. Another approach is to infer the presence of tumour-defining somatic copy number changes from shifts in average expression^1,2^. However, this approach is also subject to errors due to the choice of an appropriate baseline and expression variation not driven by somatic copy number changes. These weaknesses notwithstanding, both methods may accurately identify cancer cells in certain circumstances. However, if there is any novelty or ambiguity in the identity of cancer cells, then these two approaches are inherently fallible as they are both based on expression and not direct evidence that a cell is cancerous, i.e. that it carries the somatic cancer genome.

For example, there has been historical controversy about what cell types are malignant in neuroblastoma, a childhood cancer that arises from peripheral nervous sympathetic lineages. In addition to unambiguous cancer cells, neuroblastomas often harbour stromal cells, composed of Schwannian stroma or mesenchymal cells. It has been suggested that these stromal cell types represent cancer lineages, although a rich body of evidence, including cytogenetic investigations, have not supported this proposition^3^. Recent single cell mRNA studies of neuroblastoma have rekindled the debate on the basis of expression-based cancer cell identification^4^. Although neuroblastoma is an exemplar of the difficulties in annotating single cell tumour transcriptomes, the same problems are common to tumours with complex histology or unresolved origins. Even among tumours with well-defined origins, the variability inherent to all cancer can make annotation challenging.

The alternative to expression-based annotation is direct detection of either cancer-defining (i.e. somatic) point mutations or copy number aberrations from the nucleic acid sequences of each transcriptome, which we pursued here. Such approaches utilise additional information from whole genome/exome sequencing of tumour DNA to detect the altered genotype or the allelic imbalance it creates (**Figure 1A**).

**Figure 1.**
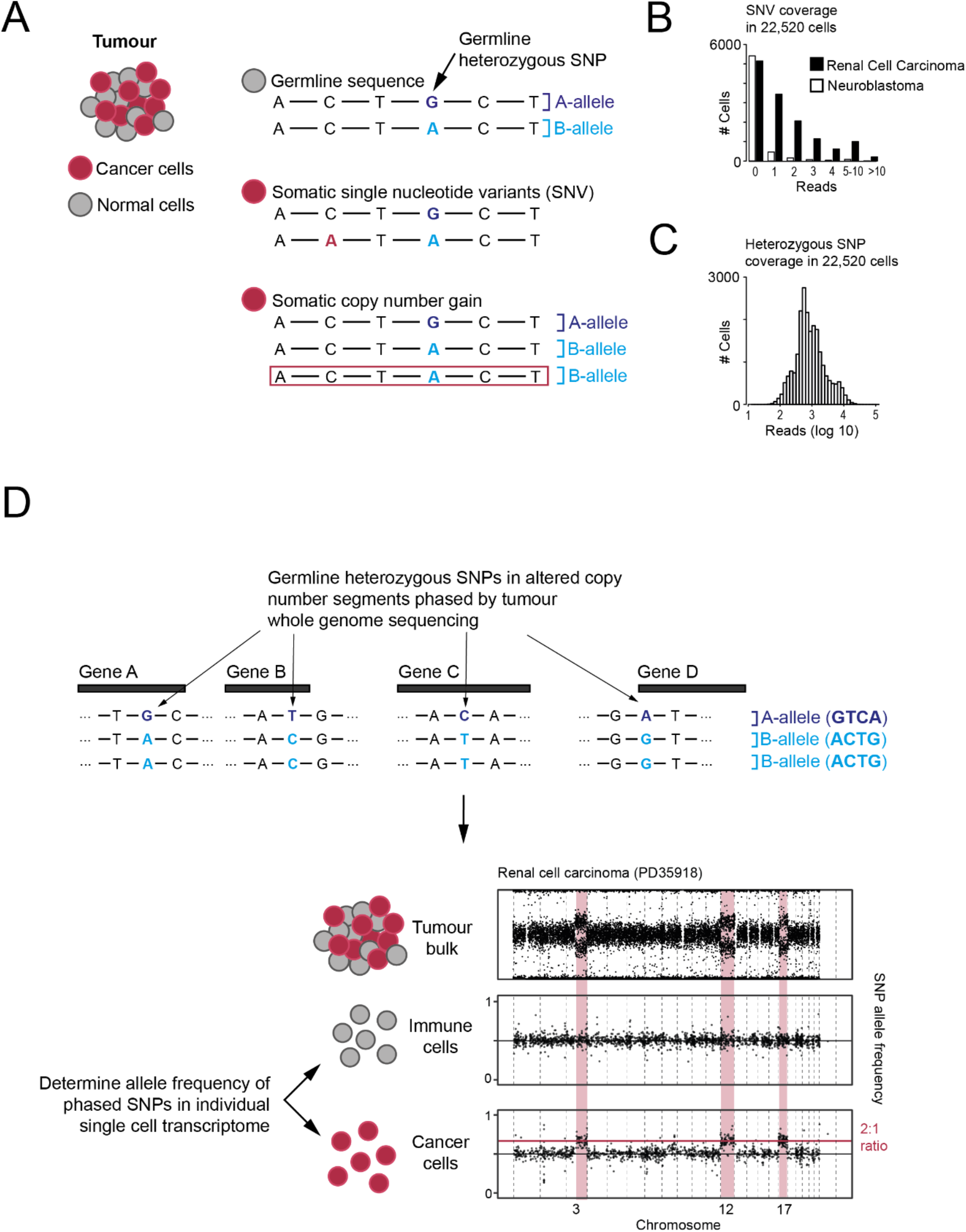
**A** - Genomic changes present in cancer genomes. **B** - Number of cells (y-axis) with N reads covering point mutations (x-axis), separated by low (NB-neuroblastoma) and high (RCC - renal cell carcinoma) mutation burden. **C** - Number of cells (y-axis) with N reads covering heterozygous single nucleotide polymorphisms (SNP)(x-axis). **D** - Overview of using allelic shifts representing copy number changes to detect cancer cells.

To test approaches used to identify cancer cells, we considered renal cell carcinoma (RCC) and neuroblastoma droplet-based 3’ single cell sequencing datasets from nine patients^5,6^. We first tested if detection of cancer specific point mutations would identify cancer transcriptomes. Across all samples, the majority of cells had no reads covering a point mutation (**Figure 1B**), with on average 3.8 reads per ten thousand point mutations per cell (range 0 to 260). This implies a tumour would need ~30,000 mutations to achieve an average coverage of 10 reads per cell, well in excess of the mutational burden of most tumours^7^. By contrast, an average of 1644 reads per cell covered heterozygous single nucleotide polymorphisms (SNPs), implying 0.6 informative reads per megabase per transcriptome (**Figure 1C**). As copy number changes may alter the allelic ratio, these data can be used to detect the cancer genotype (**Figure 1D**). This implies that a loss of heterozygosity (LoH) of 18 megabases or more should be detectable in single transcriptomes (assuming a binomial distribution and 99% accuracy).

Next, we compared the performance of cancer transcriptome identification using expression- and nucleotide-based copy number detection. For each patient we inferred the cancer genotype using a method based on shifts in average expression, CopyKAT^2^, and a statistical model based on allelic ratios^8^, which we named alleleIntegrator. We first considered RCC, an adult kidney cancer defined by expression of an unusually specific tumour marker, *CA9*, caused by the near universal disruption of the *VHL* gene underpinning RCC^9^ (**Figure 2A**). Of the 1718 cells designated as tumour in the published annotation, expression-based identification called 931 as tumour and 200 as normal, while alleleIntegrator identified 795 as tumour and 56 as normal. Concerningly, CopyKAT’s expression-based identification classed 84% (2952/3519) proximal tubular cells as cancerous. Only 0.3% (11 cells) were identified as cancer-derived by allelic ratio (**Figure 2B**). These proximal tubular cells had been derived from a macroscopically and histologically normal tissue biopsy obtained from an uninvolved region of the kidney, and did not express *CA9*. As proximal tubular cells are the probable cell of origin for RCC, it is likely that expression-based annotation incorrectly identified proximal tubular cells as cancer-derived due to their transcriptional similarity to RCC cells. Next, we assessed how well the inferred copy number profiles matched the ground truth — i.e. somatic copy number profiles obtained from whole genome sequences — at the chromosome level. For expression-based profiling, we marked any chromosome with an average absolute log expression ratio above 0.2 as significantly altered, revealing copy number aberrations (CNAs) called where none existed and true CNAs not being called (**Figure 2C, S1**). By contrast, the allelic ratio of the tumour cells and normal cells matched the ground truth (**Figure 2C, S1**). In aggregate, these analyses demonstrate the potential for expression-based methods to misidentify normal cells as cancerous, illustrating their shortcomings in identifying novel cancer cell types.

**Figure 2.**
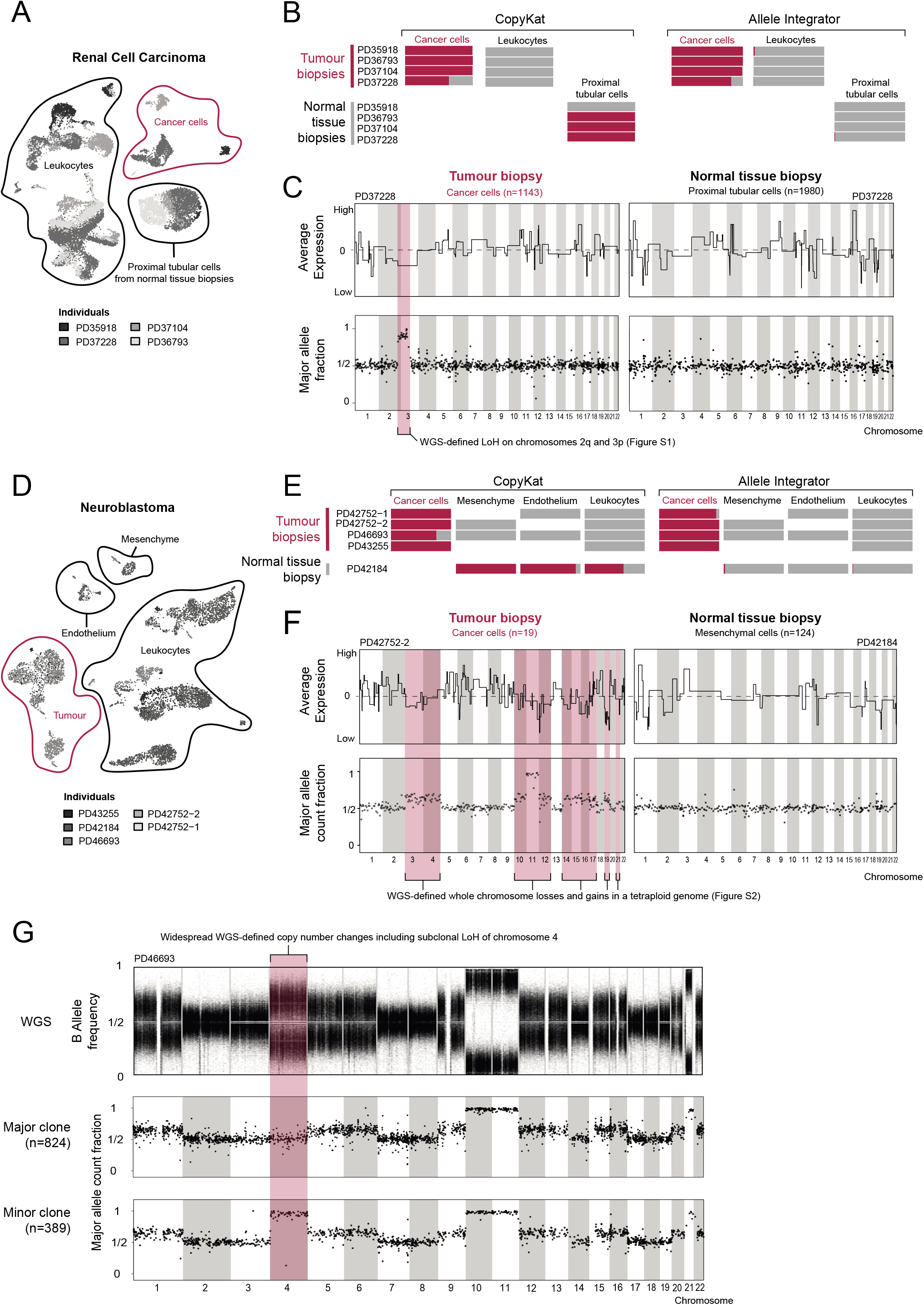
**A** - UMAP of RCC single cell transcriptomes showing patient (shading), cell type (contours and labels), and patient composition (barplots). Inset shows expression of RCC marker *CA9*. PTC - proximal tubular cells derived from normal biopsies. **B** - Cancerous (red) and non-cancerous (grey) cell fraction excluding ambiguous cells by cell type (x-axis) and sample/region (y-axis) called by CopyKAT (left) or alleleIntegrator (right). **C** - Copy number profile for PD37228 tumour (left) and proximal tubular (right) clusters from normalised averaged expression (top panels, solid black line) and allelic ratio (bottom panel, one dot per bin with ~500 reads), with true copy number changes from DNA (red shading). **D-F** - As per **A-C** but for neuroblastoma. **G** - Allelic ratios (y-axis) across the genome (x-axis) from bulk tumour DNA (top panel), cells assigned to the major clone (middle panel) and minor clone (bottom panel).

We next tested cancer transcriptome identification on single cell transcriptomes from neuroblastomas, which have no definitive single marker equivalent to *CA9* in RCC (**Figure 2D**). As before, both expression- and allelic ratio-based identification identified tumour cells accurately (**Figure 2E**). However, for sample PD42184, which is derived from a normal adrenal gland and therefore contains no tumour cells, expression-based annotation predicted 1926 cancer derived cells, including mesenchymal cells (**Figure 2E**). By contrast, these cells are identified as normal based on their allelic ratio (**Figure 2E, S2**). Similarly to the RCCs, the true copy number profile was not recovered by the average expression in any of the neuroblastomas, with an average of 4.8 false positive and 66 false negative copy number changes per sample (**Figure 2F, S2**). Of particular note, the average expression shift in mesenchymal cells is consistent with what is observed in cancer cell clusters in other samples, with significant shifts in average expression on chromosomes 1,2,3 and 12 (**Figure 2F**). Despite the complex copy number profiles of these neuroblastomas, the allelic ratios of single cell transcriptomes matched the ground truth in all cases (**Figure 2F, S2**). Overall, this demonstrates the risk of drawing erroneous biological conclusions, in this case that mesenchymal cells are cancer-derived, when relying on expression based annotation of cancer transcriptomes.

Beyond distinguishing cancer and normal cells, the high precision of copy number genotyping by allelic ratios may lend itself to the identification of minor cancer cell populations (subclones) defined by copy number aberrations. We investigated cancer subclone identification in a neuroblastoma (PD46693) that harboured a minor clone, comprising ~30% of cells, defined by copy number neutral loss of heterozygosity of chromosome 4. AlleleIntegrator identified 389/1282 subclonal cells with a posterior probability of more than 99% (**Figure 2G**). These cells are transcriptionally extremely similar, with only 95 genes and 7 transcription factors significantly differentially expressed between the major and minor clones (**Figure S3, Table S1, S2**). Amongst these genes were neuroblastoma-associated genes *NTRK1*, *BCL11A*, *TH* and *CHGB*, as well as *HMX1*, a transcription factor on chromosome 4 that is a master regulator of neural crest development. Although we would not claim that these genes collectively or individually are the definitive target of the subclonal copy number change, this analysis illustrates the power of our approach in deriving functional hypotheses about copy number changes. This is particularly pertinent in neuroblastoma, where clinical risk is defined by segmental copy number changes that remain functionally cryptic^10^.

We have shown that allelic imbalances that represent cancer-defining somatic copy number changes can precisely identify single cancer cell transcriptomes. A prerequisite of this approach, that limits its application, is the presence (and knowledge) of somatic copy number changes. We consider the main utility of our approach to lie in corroborating or refuting claims of novel cancer cell types and for investigating the functional consequences of subclonal copy number changes. Where direct nucleotide interrogation is not feasible, the expression of marker genes and detection of average shifts in expression with tools such as CopyKAT, may still provide a reasonable basis for indirectly inferring which single cell transcriptomes possess the somatic cancer genotype. However, our observations caution against identifying novel cancer cell types through such approaches alone, without direct interrogation of underlying nucleotide sequence. Accordingly, our findings suggest that it may be warranted to reappraise recent claims of novel cell types in a variety of cancers, such as neuroblastoma, that were solely based on expression based cancer cell identification.

