## Supplementary Materials for "Precise identification of cancer cells from allelic imbalances in single cell transcriptomes"

### Methods

#### Identifying cancer cells using allelic ratio

To identify cancer cell transcriptomes, we used a bayesian statistical framework[1,2], implemented in an R package, alleleIntegrator. The calling of cancer cell transcriptomes has 4 steps (**Figure 1A)**:

1. Call copy number changes and heterozygous SNPs.
2. Phase heterozygous SNPs within regions with altered copy number using tumour DNA.
3. Count reads supporting the major/minor allele in each copy number segment/transcriptome.
4. Calculate posterior probability of the cancer and normal genotype.

The precise step-by-step implementation is contained in the provided code and a detailed description of each step is provided below.

##### Calling heterozygous SNPs and copy number changes

We identified copy number (CN) states using Battenberg[3] applied to whole genome sequencing of tumour DNA. Heterozygous SNPs were called using bcftools mpileup/call to find sites with reads supporting two alleles and a B allele frequency (BAF) between 0.2 and 0.8. Sites inconsistent with heterozygosity were excluded using a binomial test with 5% FDR[4]. Alternatively, CN states and heterozygous SNP locations can be provided using alternate methods.

##### Phasing heterozygous SNPs in copy number region(s)

Using alleleCount (https://github.com/cancerit/alleleCount), we counted reads supporting each allele in tumour DNA in regions of uneven CN (i.e., where the number of maternal/paternal copies differ). The reference (or alternate) allele was assigned to the minor allele when the BAF was greater than (or less than) 0.5. Sites not significantly different from 0.5 (binomial test, 5% FDR) were excluded.

##### Counting reads by allele in each transcriptome

At each phased SNP, we calculated the counts supporting the major and minor allele for each transcriptome using alleleCount in 10X mode (-x flag). These were summed by segment/transcriptome, producing a table of major and minor allele counts for each transcriptome and copy number segment.

##### Calculating posterior probability of cancer genotype

We aimed to compare two possibilities: that the cell contains the cancer genotype or the normal genotype. To this end, we constructed a model that accounts for the major known error processes and properties of transcription: errors can alter the observed allele, transcription occurs in bursts, and transcription can exhibit allelic bias. We used a negative binomial likelihood, where the overdispersion captures extra variability due to transcriptional bursts.

We first filter out SNPs that: are imprinted (i.e., only ever express one allele), are not intronic or exonic, or have zero coverage. This filtering is most accurate when cells with the normal genotype can be specified (e.g. leukocytes in a solid tissue tumour). We also exclude genes known to display complex allele-specific expression (ASE), specifically, *HLA* and *HB* genes.

We specify a site-specific error rate of 0.01 for exonic reads and 0.05 for intronic reads, calibrated by counting non-reference reads at sites homozygous for the reference.

After filtering, we calculate the posterior probability of allele-specific expression in normal cells for each gene, using a beta distribution prior with mean 0.5 and spread set manually or to the best fit value of highly expressed genes (default genes > 400 counts). Where normal cells are not given, both alleles are considered equally likely.

Next, we calculate the maximum likelihood value of the negative binomial overdispersion from normal cells using the error rate and ASE values derived above. We optionally marginalise this estimate over the ASE posterior distribution, although we find this step makes no difference to the final estimate. Where normal calls are not given, the overdispersion is set manually or the best fit is calculated across all cells. Including non-normal cells increases the over-dispersion, making downstream calls of which cells are cancer-derived more conservative.

The expected allelic ratio at each SNP is then given by

$$\left(\frac{f\rho}{f\rho + (1-f)(1-\rho)}\right)(1-2\epsilon) + \epsilon$$

where $f$ is the number of major copies of the segment as a fraction of the total (i.e., 0.5 for diploid, ⅔ for a gain of one copy, 1 for the loss of one copy), $\boldsymbol{\rho}$ is the ASE ratio (i.e., 0.5 for no ASE) and ε is the site-specific error rate. A genome-wide posterior probability is then calculated for the cancer and normal genotype (using a flat prior) and cells are assigned as cancer or normal where this exceeds 0.99.

### Data and code availability

Previously published data was obtained for renal cell carcinoma[1] and neuroblastoma[2]. The R package, alleleIntegrator, is available from (https://github.com/constantAmateur/alleleIntegrator) and code used to generate these results from (https://github.com/mitrinh1/scGenotyping).

#### Data QC, clustering, and visualization

We used R (v4.0.4) and Seurat (v.4.0.3) for these analyses. Cells with <200 genes, <600 UMIs, mitochondrial fraction exceeding 20% (30% for RCC normal tissue), or Scrublet[5] doublet score >0.5 were excluded. High resolution clusters (resolution=10) with >50% cells failing QC were also excluded.

Data were log normalized and scaled, and principal components were calculated using highly variable genes using the standard Seurat workflow. Louvain clustering was performed with resolution 1 and a uniform manifold approximation and projection (UMAP) calculated, using 55 and 25 principal components for NB and RCC datasets respectively. Finally, cells were labelled using the published annotation and leukocytes, endothelium, mesenchyme, proximal tubular cells and tumour cells were retained.

#### Coverage of point mutations and heterozygous SNPs

For all samples, heterozygous SNPs were identified (as described above) and point mutations were called against the GRCh37d5 reference as previously described[1,2]. Coordinates were lifted over to GRCh38 and counts covering point mutations and SNPs were calculated for each transcriptome using allele counter.

#### Calling copy number aberrations

Copy number profiles and tumour subclonality was determined using Battenberg (v2.2.5). Segments shorter than 1Mb or 10% of the chromosome were removed as likely artifacts. Chromosomes were set to the same state where >=90% had a particular change and gaps filled where consecutive segments had the same copy number state and were < 1Mb apart. Subclonal copy number segments were defined as those with a second copy number state detected in a smaller fraction of tumour cells (>=10% but <50%) and are longer than 20Mb.

#### Evaluating accuracy of transcriptome classification

CopyKAT[6] (v0.1.0) was run with default parameters per-sample using cellranger filtered counts and 80% of leukocytes (randomly selected) specified as normal. This generated expression profiles on a log scale and classification (diploid, aneuploid, or uncalled) for all cells. Expression profiles were averaged by cell type and region with absolute averaged expression >0.2 in a 25Mb window were marked as altered. To visualise the allelic ratio, allele-specific counts were aggregated by cell type into bins chosen such that each bin contained at least 500 counts.

#### Analysis of PD46693 subclone

Cells with posterior probability greater than 0.99 of loss of heterozygosity of chromosome 4 in PD46693 were assigned to the subclone, those with posterior probability less than 0.01 were assigned to the major clone, and all others were called ambiguous.

Differential gene expression was performed using negative binomial regression in DESeq2[7], treating cells in the major/minor clone as replicates and removing genes with <=10 reads. We separately tested all genes and just transcription factors for significance, using a multiple hypothesis corrected[4] p-value cut-off of 0.01.

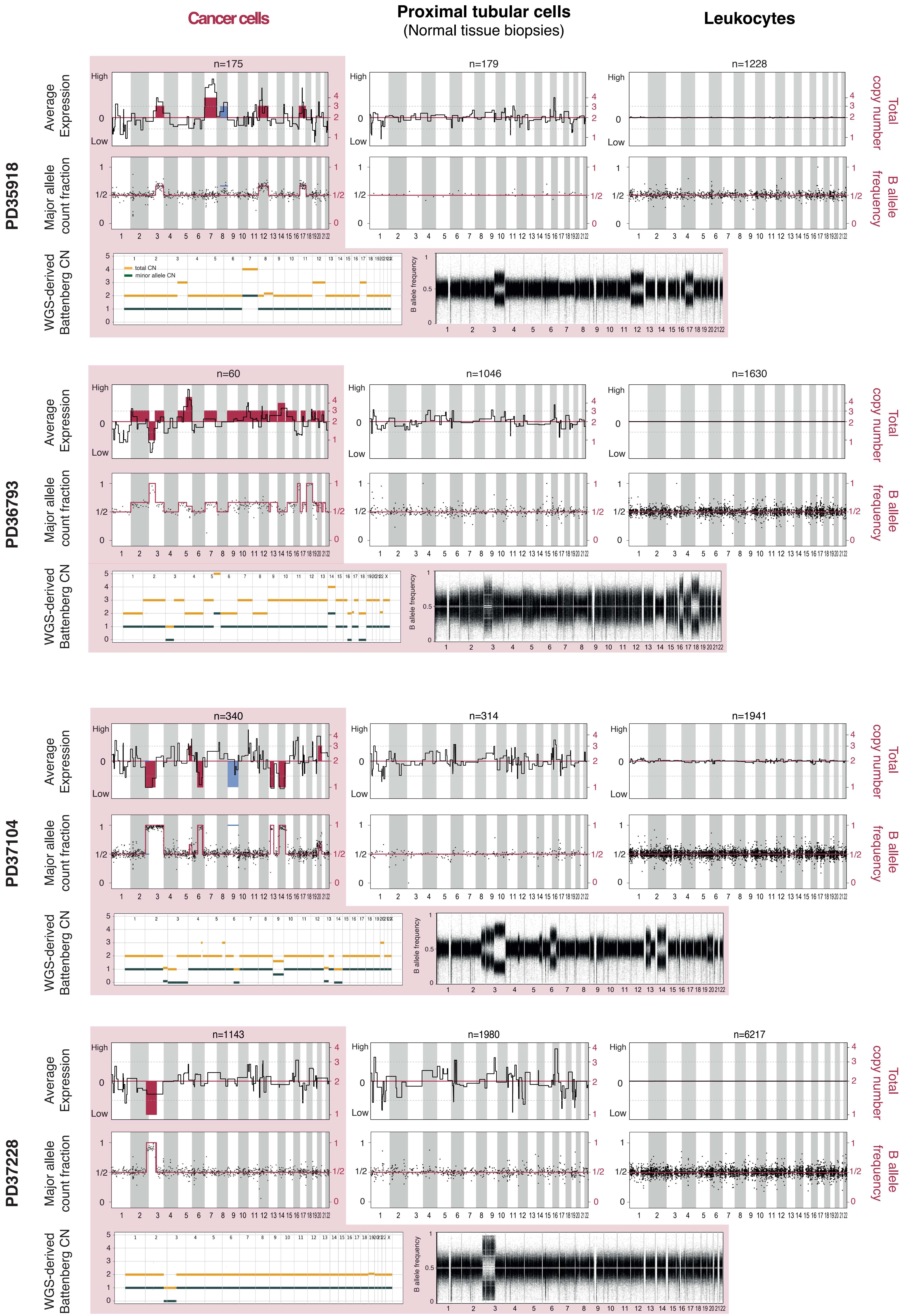
Cancer cells
Proximal tubular cells
(Normal tissue biopsies)
Leukocytes
PD35918
PD36793
PD37104
PD37228
n=175
n=179
n=1228
n=60
n=1046
n=1630
n=340
n=314
n=1941
n=1143
n=1980
n=6217
Average Expression
High
Low
Major allele count fraction
WGS-derived Battenberg CN
total CN
minor allele CN
B allele frequency
Total copy number
B allele frequency

#### Figure S1

Copy number profile for 5 renal cell carcinoma samples from normalised averaged expression (top panels, solid black line) and allelic ratio (middle panel, one dot per bin with ~500 reads), with ground truth from WGS (red, arbitrary scale in top panel). Sub-clonal copy number changes shown in blue. The dashed line (top panels) is average absolute log expression ratio of 0.2.

Bottom panel of each sample shows the WGS-defined copy number changes by Battenberg (green line represents the minor allele copy number state, yellow line represents total copy number state), and the corresponding WGS–defined B allele frequency.

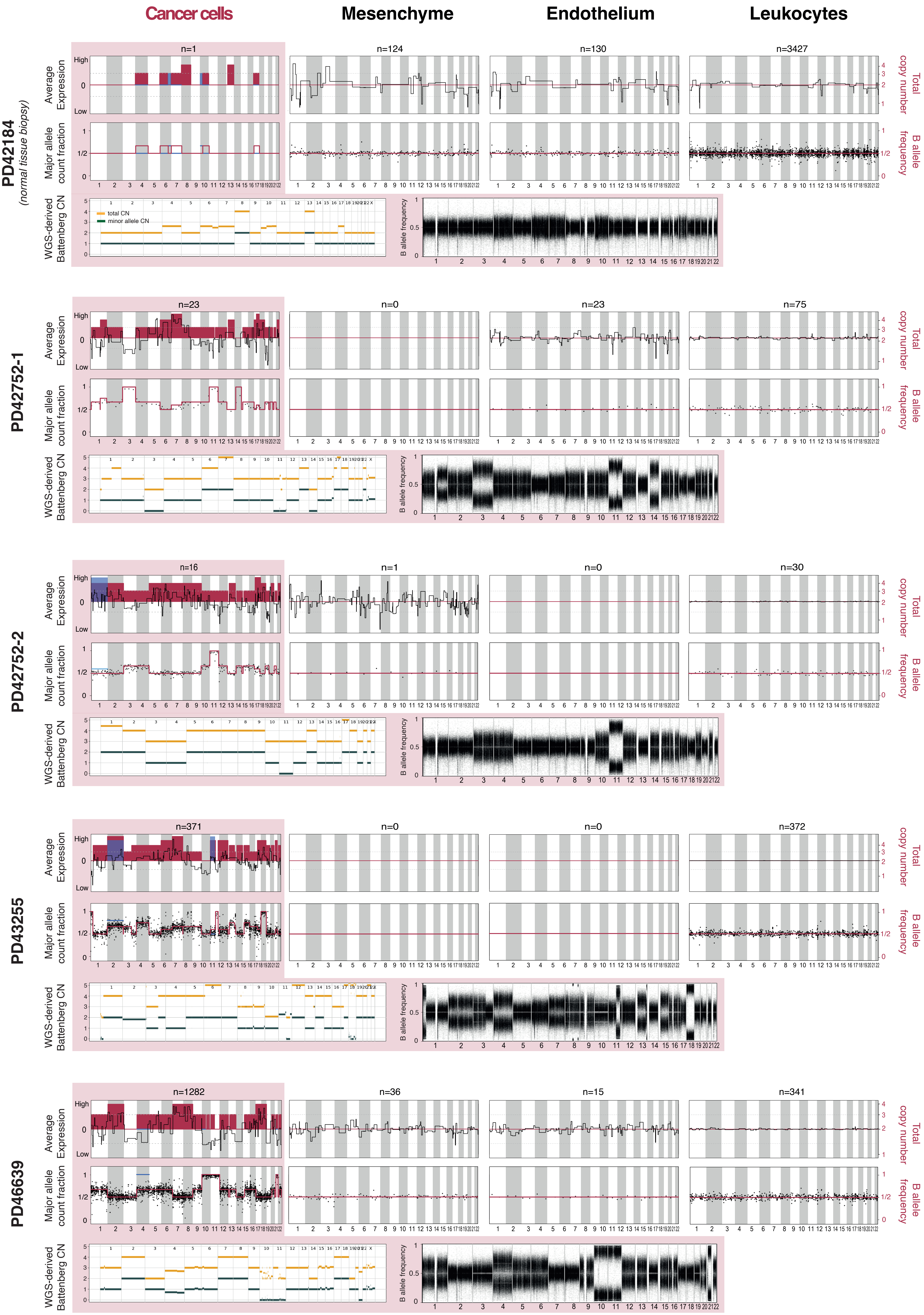
Cancer cells
Mesenchyme
Endothelium
Leukocytes
PD42184
(normal tissue biopsy)
n=1
n=124
n=130
n=3427
PD42752-1
n=23
n=0
n=23
n=75
PD42752-2
n=16
n=1
n=0
n=30
PD43255
n=371
n=0
n=0
n=372
PD46639
n=1282
n=36
n=15
n=341
Average Expression
High
Low
Major allele count fraction
1/2
WGS-derived Battenberg CN
total CN
minor allele CN
B allele frequency
Total copy number
B allele frequency

**Figure S2**

As per **Figure S1** but for neuroblastoma.

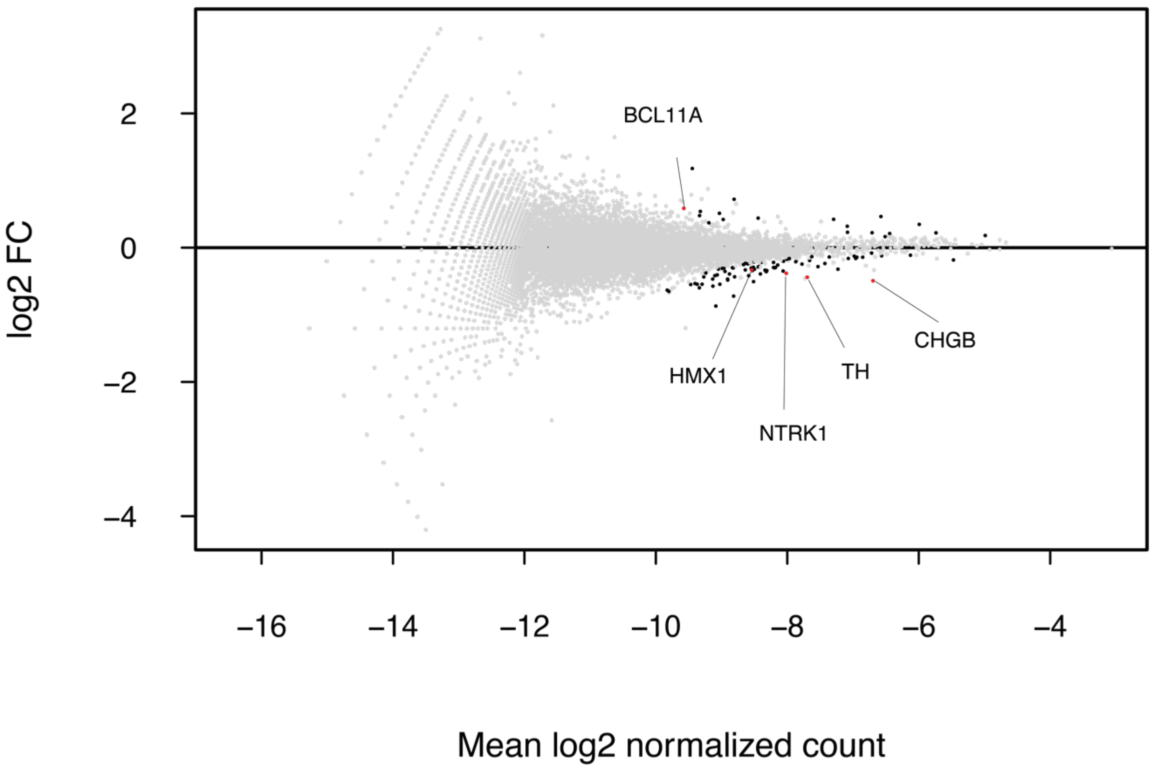


**Figure S3**

Log fold change (y-axis) for each gene (points) by average normalised expression (x-axis), with differentially expressed genes/transcription factors shown in black.

**Table S1:** List of differentially expressed genes between PD46693 minor and major clone tumour cell population.

**Table S2:** List of transcription factors that are differentially expressed between PD46693 minor and major clone tumour cell population.

**Table S3:** Copy number profile and subclonality of sample PD35918 as defined by Battenberg using WGS.

**Table S4:** Copy number profile and subclonality of sample PD36793 as defined by Battenberg using WGS.

**Table S5:** Copy number profile and subclonality of sample PD37104 as defined by Battenberg using WGS.

**Table S6:** Copy number profile and subclonality of sample PD37228 as defined by Battenberg using WGS.

**Table S7:** Copy number profile and subclonality of sample PD42184 as defined by Battenberg using WGS.

**Table S8:** Copy number profile and subclonality of sample PD42752-1 as defined by Battenberg using WGS.

**Table S9:** Copy number profile and subclonality of sample PD42752-2 as defined by Battenberg using WGS.

**Table S10:** Copy number profile and subclonality of sample PD43255 as defined by Battenberg using WGS.

**Table S11:** Copy number profile and subclonality of sample PD46693 as defined by Battenberg using WGS.
